# Simultaneous assessment of mechanical and electrical function in Langendorff-perfused ex-vivo mouse heart

**DOI:** 10.1101/2023.09.15.557526

**Authors:** Julien Louradour, Rahel Ottersberg, Adrian Segiser, Agnieszka Olejnik, Berenice Martínez-Salazar, Mark Siegrist, Manuel Egle, Miriam Barbieri, Saranda Nimani, Nicolò Alerni, Yvonne Döring, Katja E. Odening, Sarah Longnus

## Abstract

**Background:** The Langendorff-perfused *ex-vivo* isolated heart model has been extensively used to study cardiac function for many years. However, electrical and mechanical function are often studied separately - despite growing proof of a complex electro-mechanical interaction in cardiac physiology and pathology. Therefore, we developed an isolated mouse heart perfusion system that allows simultaneous recording of electrical and mechanical function.

**Methods:** Isolated mouse hearts were mounted on a Langendorff setup and electrical function was assessed via a pseudo-ECG and an octapolar catheter inserted in the right atrium and ventricle. Mechanical function was simultaneously assessed via a balloon inserted into the left ventricle coupled with pressure determination. Hearts were then submitted to an ischemia-reperfusion protocol.

**Results:** At baseline, heart rate, PR and QT intervals, intra-atrial and intra-ventricular conduction times, as well as ventricular effective refractory period, could be measured as parameters of cardiac electrical function. Left ventricular developed pressure (DP), left ventricular work (DP-heart rate product) and maximal velocities of contraction and relaxation were used to assess cardiac mechanical function. Cardiac arrhythmias were observed with episodes of bigeminy during which DP was significantly increased compared to that of sinus rhythm episodes. In addition, the extrasystole-triggered contraction was only 50% of that of sinus rhythm, recapitulating the “pulse deficit” phenomenon observed in bigeminy patients. After ischemia, the mechanical function significantly decreased and slowly recovered during reperfusion while most of the electrical parameters remained unchanged. Finally, the same electro-mechanical interaction during episodes of bigeminy at baseline was observed during reperfusion.

**Conclusion:** Our modified Langendorff setup allows simultaneous recording of electrical and mechanical function on a beat-to-beat scale and can be used to study electro-mechanical interaction in isolated mouse hearts.

## 1 Introduction

Since its development in the 19th century, the Langendorff isolated heart model has made a crucial contribution to our understanding of cardiac pathophysiology and remains to this day a valuable tool for studying the mechanical, electrophysiological, metabolic, vascular and biochemical processes of the heart in a variety of disease settings and animal models (1–3). With the isolated heart system, all of these mechanisms can be investigated in the intact organ, preserving the multiple, interdependent cell types and function, but without the potential confounding influence of systemic effects. Furthermore, the experimental conditions can be tightly controlled which leads to highly reproducible results (4). One frequent use for the isolated heart model is to investigate strategies for cardioprotection in the context of ischemia-reperfusion injury (IR) (5). Indeed, coronary heart diseases are the leading cause of death worldwide, significantly contribute to the global burden of disease (6–8), and are associated with a variety of potentially fatal complications, a common one being IR-induced arrhythmias (9).

Isolated mouse heart models have been widely used to test the effect of gene mutations on cardiac physiology due to the fast and easy manipulation of the mouse genome. However, in most of the isolated mouse heart models used to investigate IR described in scientific literature, contractile and electrical cardiac function are studied separately. Yet, there is a complex relationship between electrical and mechanical function in cardiac physiology. Electrical excitation triggers the contractile activity of the heart (through excitation-contraction coupling), but mechanical activity can also affect the electrical activity (through mechano-electrical feedback) (10–12). Indeed, changes in cardiac pressure and/or volume can affect cardiac depolarization or repolarization and may trigger arrhythmias (10,13,14). Given this strong mutual influence of excitation-contraction coupling and mechano-electrical feedback and their involvement in pathophysiology (10), there is a growing need for new approaches to study this electro-mechanical interaction in general, but also in specific, pathophysiological settings such as ischemia-reperfusion.

Therefore, we developed an isolated mouse heart perfusion system to simultaneously assess cardiac electrical and mechanical function. We showed that our model can recapitulate pathophysiological electro-mechanical interactions, and we tested this approach in a specific example of a global ischemia-reperfusion protocol. We finally suggest that this approach might be useful to better understand the complex mutual influence between electrical and mechanical cardiac function.

## 2 Materials and equipment

Equipment and reagents used in this study are listed in Tables 1 and 2.

**Table 1.** List of equipment.

| Perfusion system |  |  |
| --- | --- | --- |
| Component | Description and function | Supplier |
| 1.1F Intracardiac octapolar catheter | Placed in the right atrium and ventricle to record the electrical signal with 8 equally interspaced electrodes | EPR-800, Millar Instruments, Houston, Texas, United States |
| Aortic pressure transducer | Situated in the aortic line and provides feedback for control of pump speed | SP 844, MEMSCAP, Crolles, France |
| Compliance chamber | Works as a compliance element as well as bubble trap | Radnoti LTD, Dublin, Ireland |
| Double walled tubing | Double-walled tubing to ensure a suitable buffer temperature with the circulation of warm water through the walls of the tubes | Radnoti LTD, Dublin, Ireland |
| In-line flowmeter | Situated downstream of peristaltic pump to measure coronary flow | ME3PXN, Transonic Systems Inc., Ithaca NY, United States |
| Intraventricular silicone balloon | Silicone balloon inserted into the left ventricle and connected to a pressure transducer to measure left ventricular pressures | Custom made of ALPA-Sil 32 silicone (40-2383A/B, Silitec AG, Gümligen, Switzerland) |
| Ischemic bath | Bath filled with warm (37°C), energy-substrate-free KHB gassed with 95% N<sub>2</sub> and 5% CO<sub>2</sub> used to immerse the heart during warm ischemia | Radnoti LTD, Dublin, Ireland |
| LabChart pro v8 | Data analysis software to record and display all the acquired signals | AD Instruments Ltd, Oxford, United Kingdom |
| Langendorff reservoir with glass tube gas disperser | Reservoir to collect and oxygenate buffer from the non-recirculating line | Radnoti LTD, Dublin, Ireland |
| Millar catheter | Connected to the intraventricular balloon to measure left ventricular pressure | SPR-671NR, Millar Inc., Houston, USA |
| Mini oxygenator | Low-volume membrane oxygenator to oxygenate the buffer from the recirculating line | Custom built |
| Myopacer stimulator | Required to pace the ventricles with a S1-S2 protocol to determine the ventricular effective refractory period | MyoPacer stimulator, IonOptix, Westwood, Massachusetts |
| Peristaltic pump | Drives both circuits (recirculating and non-recirculating), controlled by a feedback mechanism (STH Pump controller) using a pressure sensor in the aortic line to perfuse the heart at a constant pressure (~60mmHg) | Minipuls 3, Gilson Incorporated, Middleton, Wisconsin, USA |
| Powerlab | Data acquisition instrument used to record all perfusion data | AD Instruments Ltd, Oxford, United Kingdom |
| Pump controller | Translate the pressure feedback from the aortic line into an analog signal to control the speed of the peristaltic pump | STH pump controller, AD Instruments Ltd, Oxford, United Kingdom |
| Recirculating reservoir | Collects the buffer that comes from the heart and feeds it back into the circulation | Radnoti LTD, Dublin, Ireland |
| Stereomicroscope | To cannulate the mouse heart | ST, s/n 220017, Optech, Munich, Germany |
| Temperature sensor | Monitors the temperature of the buffer | IT-18, Physitemp Instruments, LLC, Clifton NJ, United States |
| Ventricular pressure transducer | To measure left ventricular pressure | MLT0670, AD Instruments Ltd, Oxford, United Kingdom |

**Table 2.** List of reagents.

| Buffer |  |
| --- | --- |
| 10x Krebs-Henseleit Stock Solution |  |
| Component | Art. no., supplier |
| NaCl | 1.06404.100 Merck, Darmstadt, Germany |
| KCl | 6781.3, Roth, Karlsruhe, Germany |
| KH<sub>2</sub>PO<sub>4</sub> | 3904.2, Roth, Karlsruhe, Germany |
| CaCl<sub>2</sub> | 5239.2, Roth, Karlsruhe, Germany |
| MgSO<sub>4</sub> | P027.1, Roth, Karlsruhe, Germany |
| 1x Krebs-Henseleit Buffer (KHB) |  |
| Component | Supplier |
| (D+)Glucose | X997.1, Roth, Karlsruhe, Germany |
| NaHCO<sub>3</sub> | 6885.2, Roth, Karlsruhe, Germany |
| Pyruvate | P8574, Sigma-Aldrich, Steinheim, Germany |
| High Fat Albumin Buffer |  |
| Component | Supplier |
| Albumin | A3803-50G, Sigma-Aldrich, Steinheim, Germany |
| Ethanol | 1.00983.2511, Merck, Darmstadt, Germany |
| Palmitate | P0500-10G, Sigma-Aldrich, St. Louis, USA |
| Na<sub>2</sub>CO<sub>3</sub> | A135.1, Roth, Karlsruhe, Germany |
| Lactate | 71718-10G, Sigma-Aldrich, Steinheim, Germany |

## 3 Methods

### 3.1 Animals

All performed experiments were in accordance with EU legislation (directive 2010/63/EU) and the Swiss animal welfare law and have been approved by the Swiss animal welfare authorities (Kanton Bern, Amt für Veterinärwesen, approval number BE62-21).

Low-density lipoprotein receptor knock-out mice (LDLR^-/-^) and transgenic K18-hACE2 mice were purchased from the Jackson laboratory and cross bred to obtain K18-hACE2-Ldlr-/-mice. Eight-week-old female and male

Animals were housed in groups at 22 °C with a 12-h day/night cycle, food/water supply *ad libitum* and 12 weeks of Western-type diet (WD) to induce a cardiovascular disease phenotype. One week before the end of the WD feeding period, mice were also treated with 50 µg CpG oligodeoxynucleotides + 100 µg Polyinosinic:polycytidylic acid (Poly(I:C)) antigens every second day and three times in total to induce an inflammatory setting. The animals were euthanized three days after the last injection. These mice represent the control group of larger set of animals investigated during a Covid-19 research project, which will be published separately. However, to establish and validate the method of simultaneous mechanical and electrical function determination in Langendorff-perfused mouse hearts, as well as to allow proper understanding and replication of our experimental set up, we aimed to generate a separate paper to concentrate on technical aspects of the experimental system.

### 3.2 Langendorff setup

As shown in Table 1 and Figure 1A, the Langendorff perfusion system consisted of two separate circuits (one recirculating and one non-recirculating) converging at the aortic line. Both circuits were driven by a peristaltic pump, which was controlled via pressure feedback using a pressure transducer located in the aortic line near the cannula. All buffers were warmed to maintain the heart temperature at 37°C using a circulating water bath. Buffer oxygenation was achieved with a typical reservoir and a glass tube gas disperser in the non-recirculating circuit, while a purpose-built, low-volume membrane oxygenator similar to a previously described one (15) was used in the recirculating circuit. Perfusion buffer was oxygenated with 95% Oand 5% COduring aerobic periods and immersion buffer was gassed with 95% Nand 5% COduring ischemia.

**Figure 1.**
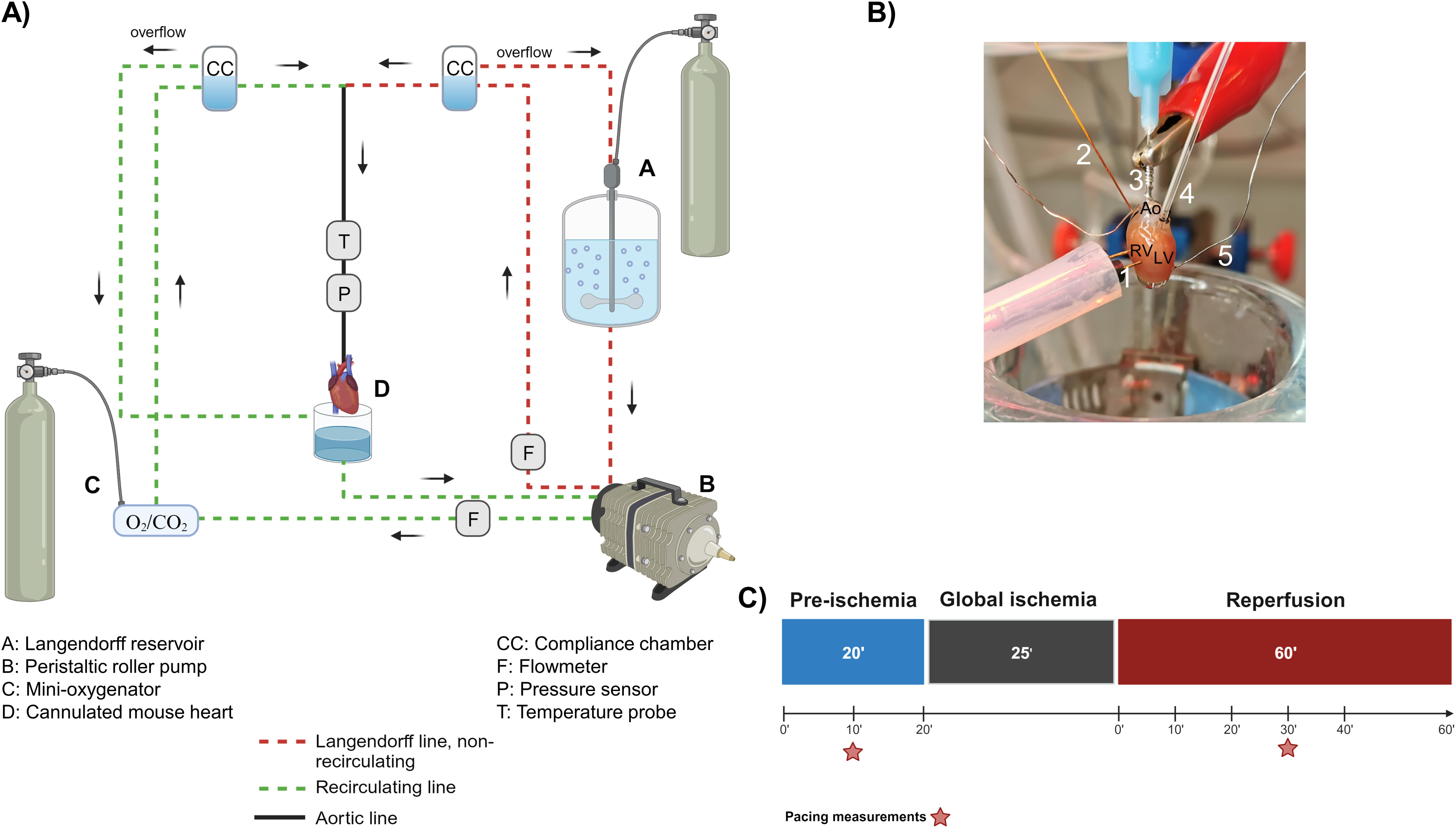
The Langendorff system. (A) General diagram of the Langendorff system and its different components. (B) Photo of a cannulated heart mounted on the Langendorff system and perfused via the aorta. The intracardiac octapolar catheter is inserted in the right atrium and right ventricle, while the balloon is inserted in the left ventricle after removal of the left atrium. 1 = pacing electrodes, 2 = intracardiac octapolar catheter, 3 = aortic cannula, 4 = intraventricular balloon system coupled to pressure measurement, 5 = silver wires, Ao = aorta, RV = right ventricle, LV = left ventricle. (C) Ischemia-reperfusion protocol during which the hearts are perfused aerobically for 20 min (pre-ischemia) before the 25 min global ischemia phase, during which perfusion is stopped, and the hearts are placed in an energy-substrate-free immersion bath. Hearts are then reperfused aerobically for 60 min. Red stars indicate the time points at which the ventricular effective refractory period was assessed by electrical pacing. Figure 1A *and 1C were created with BioRender.com*.

As illustrated in Figure 1B, a 1.1F intracardiac octapolar catheter (EPR-800, Millar Instruments, Houston, Texas) and two silver electrodes were used to measure (intracardiac and extracardiac) electrical signals. The intracardiac octapolar catheter was inserted into the right ventricle (allowing to record intraventriular and intraatrial electrical signals) and the two thin silver electrodes were placed at the root of the aorta and at the apex of the heart. For mechanical heart function, a custom-made balloon of ALPA-Sil 32 silicone (40-2383A/B, Silitec AG, Gümligen, Switzerland) was inserted into the left ventricle and coupled to either a 1.4F Millar catheter (SPR-671NR, Millar Inc., Houston, USA) or a pressure transducer (MLT0670, AD Instruments Ltd, Oxford, United Kingdom) to measure left ventricular pressure. Coronary flow was measured using two inline flowmeters (ME3PXN, Transonic Systems Inc., Ithaca NY, United States).

### 3.3 Study design and perfusion protocol

To measure the electrical and mechanical function simultaneously, isolated hearts from both male and female mice were investigated using a modified Langendorff perfusion system. All hearts were subjected to the same protocol: 20’ aerobic (baseline) perfusion, followed by 25’ normothermic, global ischemia, and 60’ aerobic reperfusion (Figure 1C).

#### 3.3.1 Heart procurement and cannulation

Mice were administered 100µg/g bodyweight ketamine (Ketalar 50mg/ml, Pfizer, Zürich, Swizerland) and 8µg/g bodyweight xylazine (Xylazin 20mg/ml, Streuli, Uznach, Switzerland) by intraperitoneal injection. The pedal pain withdrawal reflex was used to ensure sufficient depth of anesthesia. Mice were brought into a supine position and the thoracic cavity was accessed via midline skin incision and bilateral thoracotomy. The heart was then carefully explanted and transferred to a petri dish containing 50mL ice-cold 0.9% NaCl and 0.5mL of heparin (Liquemin 25000 UI/5ml, Drossopharm AG, Basel, Switzerland). While fully submerged in the solution, the aorta was cannulated with a blunt 22G needle using a stereomicroscope (ST, s/n 220017, Optech, Munich, Germany). The cannulated heart was then quickly connected to the perfusion system. The 1.1F intracardiac octapolar catheter was inserted into the right ventricle through a small incision in the right atrium and two silver wires were put in place at the root of the aorta and base of the heart. Proper positioning of the electrodes was visualized by an overlay of the intracardiac electrocardiogram signals with the P wave and QRS complex of the extracardiac electrocardiogram. Representative traces are shown in Figure 2A. After that, the left atrial appendage was removed, and a silicone balloon was inserted into the left ventricle via the mitral valve. The balloon was filled with water using a 50 µL Hamilton syringe to reach a left ventricular end-diastolic pressure (LVEDP) of ∼7-9 mmHg.

**Figure 2.**
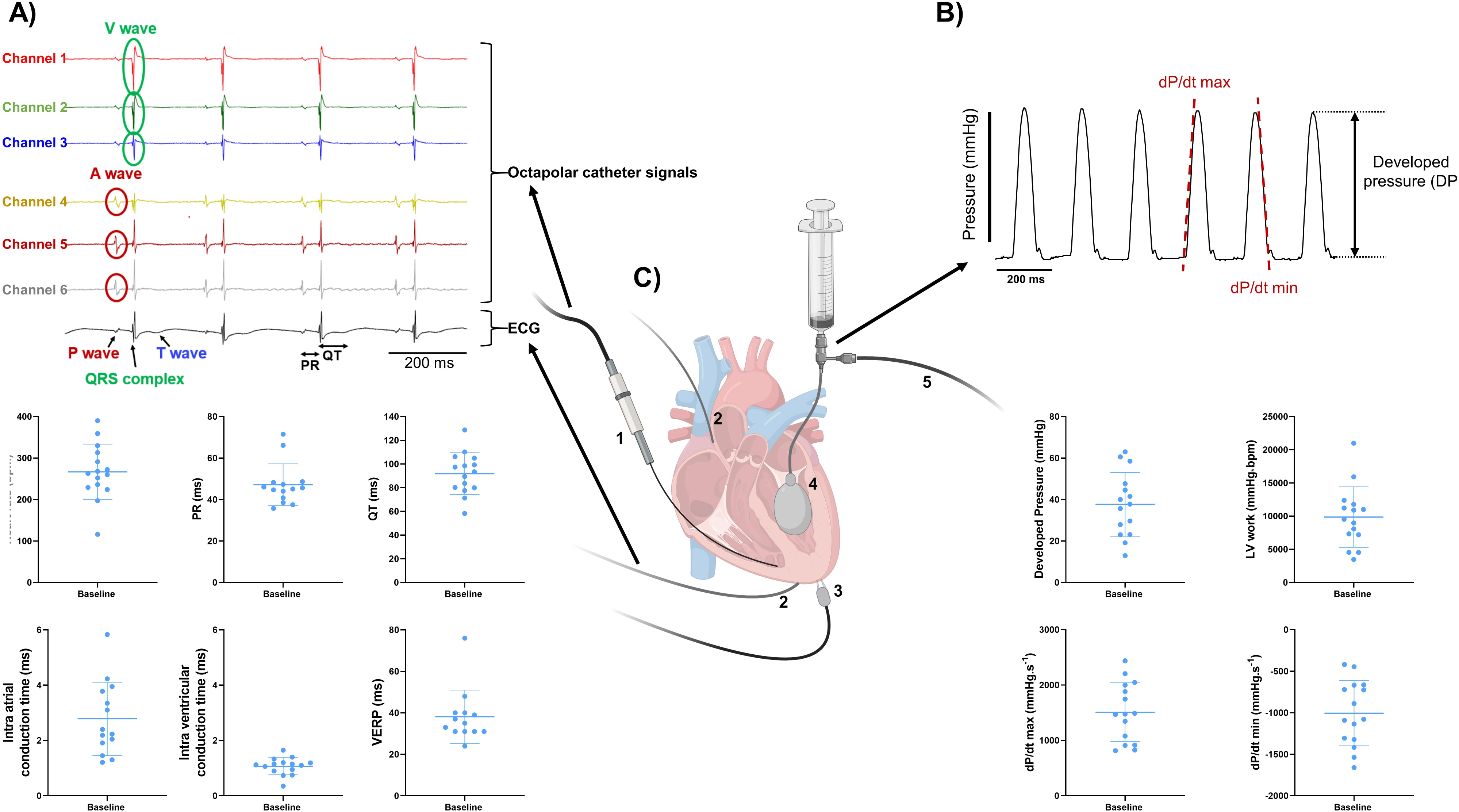
Combined measurements of electrical and mechanical function on *ex-vivo* perfused mouse hearts. (A) Example trace recordings obtained from the intracardiac octapolar catheter (channel 1-6) and ECG recording with associated average heart rate, PR interval, QT interval, intra-atrial and intra-ventricular conduction times, and ventricular effective refractory period (VERP) at baseline from the *ex-vivo* perfused hearts (N=15). Data are expressed as mean ± standard deviation (SD). (B) Example trace recording of the left ventricular pressure and associated average developed pressure (DP), left ventricular (LV) work and maximal velocity of contraction (dP/dt max) and relaxation (dP/dt min) at baseline from the *ex-vivo* perfused hearts (N=15). Data are expressed as mean ± standard deviation (SD). (C) Schematic illustration of a Langendorff-perfused heart with the intracardiac octapolar catheter inserted in the right atrium and ventricle (1), silver wires to record the electrocardiogram (2), pacing electrodes to assess the VERP (3), and the balloon (4) inserted in the left ventricle coupled to pressure recording (5) to measure left ventricular pressure. Figure 2C *was created with BioRender.com*.

#### 3.3.2 Perfusion protocol

Once connected to the system, the heart was perfused via a retrograde perfusion of the aorta with modified Krebs-Henseleit buffer (in [mM]: NaCl 116.5, KCl 4.7, KHPO1.2, CaCl1.5, MgSO1.2, (D+) Glucose 11.1, NaHCO_3_25, Pyruvate 2) using the non-recirculating Langendorff circuit at a constant pressure of 60 mmHg. The system was then switched to the recirculating Langendorff circuit for all aerobic periods and perfused with the same modified Krebs-Henseleit buffer supplemented with 1.2mM palmitate and 3% albumin at a constant pressure of 60 mmHg. During ischemia, hearts were immersed in a warm (37°Cischemic bath with energy-substrate-free modified Krebs-Henseleit buffer (in [mM]: NaCl 116.5, KCl 4.7, KHPO1.2, CaCl1.5, MgSO1.2, NaHCO25) (Table 2).

A PowerLab data acquisition system (AD Instruments Ltd, Oxford, United Kingdom) was used to monitor and record temperature, electrical signals, left ventricular pressure, and flow data throughout the entire perfusion protocol. Two-minute time windows were then analyzed after 20 minutes of baseline and 10, 20, 30, 40, and 60 minutes of reperfusion using LabChart Pro v8 (AD Instruments Ltd, Oxford, United Kingdom) as shown in Figure 1C.

#### 3.3.3 LV and electrical function measurements

Left ventricular systolic pressure (LVSP) as well as left ventricular diastolic pressure (LVDP) were measured (as shown in Figure 2). From these parameters, developed pressure (DP), left ventricular work (LV work = developed pressure * heart rate) as well as the maximal velocity of contraction (dP/dt max) and relaxation (dP/dt min) were calculated using LabChart Pro v8 (AD Instruments Ltd, Oxford, United Kingdom).

One-lead pseudo-ECG was recorded using the thin silver electrodes and later analyzed using LabChart Pro v8 (AD Instruments Ltd, Oxford, United Kingdom) to measure heart rate, PR and QT intervals. Intra-cardiac EP signals and intra-atrial and intra-ventricular conduction times were determined using the 1.1F octapolar intracardiac catheter as shown in the representative traces in Figure 2A. Intra-atrial and intra-ventricular conduction times were obtained by measuring the time delay between the first derivative negative peak from different electrodes. Intra-atrial and intra-ventricular conduction times were measured between channels 4 and 5, and 1 and 2, respectively. Finally, ventricular effective refractory period (VERP) was assessed by pacing the ventricles with a Myopacer field stimulator (IonOptix, Westwood, Massachusetts) at 10 minutes baseline and 30 minutes reperfusion (Figure 1C). S1-S2 stimulation protocol was applied with a train of 7 S1 stimulations (1ms square pulses, 5V) at a constant cycle length of 100ms, followed by one S2 stimulation with a decreasing S1-S2 interval of 1ms, starting from 100ms to 10ms. VERP was determined as the first S1-S2 interval, at which no electrical response could be triggered.

### 3.4 Data analyses

Statistical analyses were performed using Prism 9.5.1 (GraphPad Software). Data are expressed as mean ± the standard error of the mean (SEM) unless stated otherwise (Figure 2: mean ± standard deviation (SD)). Normality of the variables was assessed with Shapiro-Wilk test and parametric or non-parametric tests were used accordingly. P < 0.05 was considered statistically significant and tests used in each experiment are specified in the figure legends.

## 4 Results

### 4.1 Combined measurements of mechanical and electrophysiological function of Langendorff-perfused hearts

Our experimental setup allows a simultaneous recording of both electrophysiological and mechanical function in spontaneously beating, *ex-vivo* perfused mouse hearts. From the octapolar catheter inserted in the right atrium and ventricle (Figure 1B, 2C), the propagation of the electrical signal throughout the heart can be measured. The regularly interspaced electrodes of the EP catheter record the voltage changes over time, leading to atrial (A wave, channels 4-6) and ventricular (V wave, channels 1-3) waves (Figure 2A). By measuring the time delay between waves in consecutive electrodes in the right atrium and in the right ventricle, we can determine the atrial and ventricular conduction times, respectively (see material and method) (Figure 2A). In addition, two silver electrodes, positioned at the root of the aorta and at the apex (Figure 1B, 2C), record a pseudo-ECG, from which heart rate (HR), PR and QT intervals can be measured (Figure 2A). It is worth mentioning here that the A and V waves of the octapolar catheter signals perfectly aligned with the P wave and QRS complex of the ECG, confirming correct positioning of the electrodes in the right atrium (channels 4 to 6) and ventricle (channels 1 to 3). Finally, the ventricular effective refractory period (VERP) can be measured by pacing the ventricles with a S1-S2 stimulation protocol (Figure 1B, 2A, 2C). Combined, these recordings allow a thorough assessment of the electrophysiological function of the *ex-vivo* perfused mouse heart.

Simultaneous with the electrical recordings, mechanical function can be assessed in the same mouse heart with either a pressure transducer or Millar catheter linked to a silicone balloon inserted into the left ventricle (Figure 1B, 2C). From the left ventricular pressure curve recordings, the following mechanical function parameters can be derived: developed pressure (DP: left ventricular maximal pressure – minimal pressure), left ventricular (LV) work (HR*DP), and maximal velocity of contraction (dP/dt max) and relaxation (dP/dt min) (Figure 2B).

We tested these simultaneous measurements of electrical and mechanical activities on *ex-vivo* perfused hearts from 15 mice (7 males and 8 females) (Table 3). At baseline, hearts showed high variability and wide distribution in almost all measured parameters, as visualized with the high standard deviation (SD) bars (Figure 2). On average, hearts had a frequency of 267 ± 67 bpm, a PR interval of 47.1 ± 10.1 ms, a QT interval of 91.9 ± 17.5 ms, an intra-atrial and intra-ventricular conduction times of 2.78 ± 1.3 and 1.07 ± 0.3 ms, and a VERP of 38 ± 13 ms, expressed as mean ± SD. As for the mechanical function, DP was 37.7 ± 15.4 mmHg, LV work was 9851 ± 4548 mmHg*bpm, and dP/dt max and min were, respectively,1509.8 ± 531.0 and -1005.5 ± 393.0 mmHg.s^-1^, expressed as mean ± SD. We then examined whether electrical and mechanical parameters correlated with the heart rate (Supplementary Figure 1). As expected, QT interval was negatively correlated to heart rate (Spearman coefficient r = -0.7464, p-value = 0.002, Supplementary Figure 1A), e.g., shorter at higher heart rates, and coronary flow was positively correlated to heart rate (r = 0.6143 p-value = 0.017, Supplementary Figure 1B). However, there was no significant correlation between developed pressure and heart rate (Supplementary Figure 1C) or between dP/dt min and QT interval (Supplementary Figure 1D).

**Table 3.**
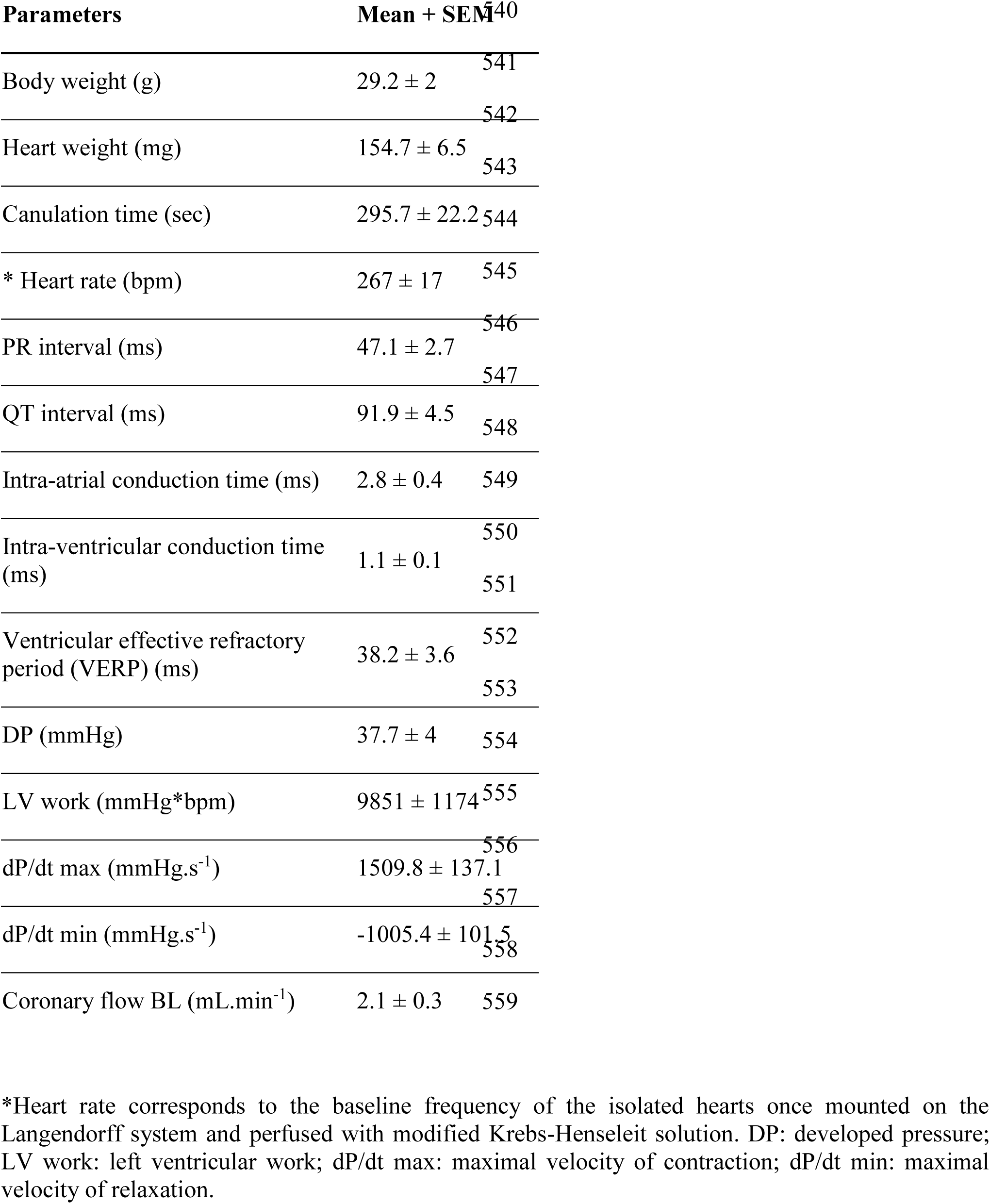
Baseline electrical and mechanical measured parameters.

### 4.2 Decreased mechanical function during spontaneous episodes of bigeminy

The advantage of simultaneous assessment of electrical and mechanical function is that it allows a better understanding of the electro-mechanical interaction, i.e., how electrical activity impacts on contraction, and vice versa. We, therefore, looked at spontaneous arrhythmia occurrence in *ex-vivo* perfused hearts to see how the mechanical function reacts to a change in the electrical activity.

The combined use of the intracardiac octapolar catheter with the ECG allowed us to better define arrhythmia, such as extrasystole, and whether the ectopic signal originates from the atria (premature atrial contraction, PAC) or the ventricles (premature ventricular contraction, PVC) (Supplementary Figure 2A-B). At baseline, hearts were either presenting with normal sinus rhythm, AV block or intermittent bigeminy (premature beat every two beats) (Figure 3A-C). Of the 15 hearts, 8 were in sinus rhythm (SR), 2 showed sustained 1^st^ degree and 2^nd^ degree AV blocks, while 5 had short episodes of bigeminy (Figure 3D, E). Notably, isolated premature beats were also observed in 3 of the 8 sinus rhythm hearts, accounting for less than 10% of the total number of beats (Supplementary Figure 2C).

**Figure 3.**
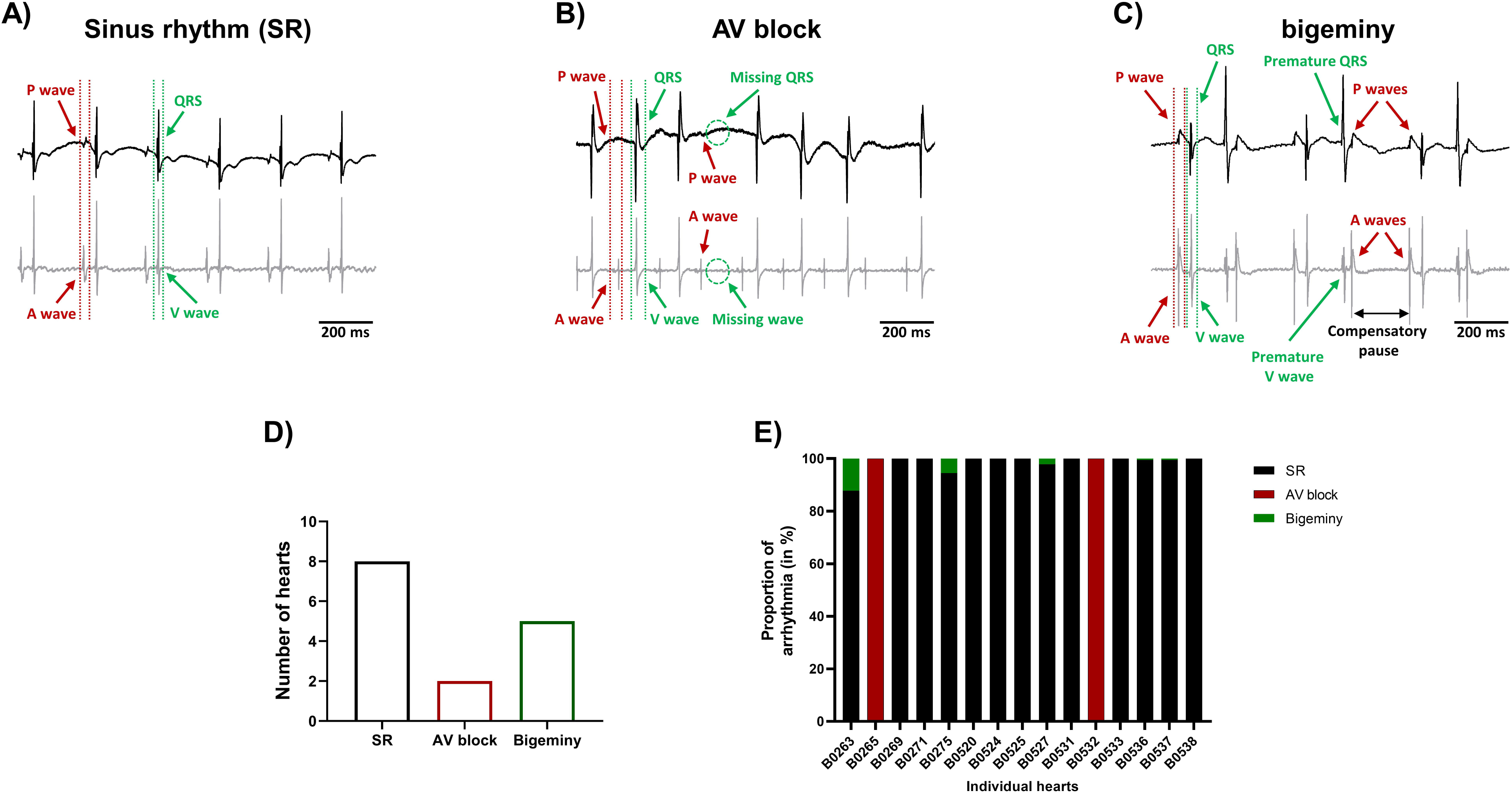
Spontaneous arrhythmia at baseline. Example recordings of the electrocardiogram trace (top, black) and one signal trace (channel 6, see Figure 2) from the octapolar catheter (bottom, gray) of sinus rhythm (SR) (A), AV block (B) and bigeminy (C) episodes at baseline. Red and green arrows indicate atrial and ventricular signals, respectively. (D) Proportion of *ex-vivo* hearts showing sinus rhythm, AV block or bigeminy at baseline (N=15). (E) Proportion of arrhythmia occurrence during the considered baseline time-window in individual hearts. Black = SR, red = AV block and green = bigeminy.

To see how electrical dysfunction might impact the mechanical function, we compared the contractility of the left ventricle during episodes of bigeminy in hearts presenting with this arrhythmia at baseline to that of sinus rhythm hearts. Though not significant, there were increases in average DP and dP/dt max, as well as decreases in LV work and dP/dt min when comparing hearts with bigeminy versus sinus rhythm (Figure 4A-D). Since the bigeminy episodes were intermittent and accounted for less than 20% of the total time period (Figure 3E), we could compare the left ventricular pressure during episodes of bigeminy and episodes of sinus rhythm within the same hearts. This intracardiac comparison showed a significant increase in DP (SR: 31.9 ± 3.3; bigeminy: 46.0 ± 7.9 mmHg) and dP/dt max (SR: 1375.6 ± 216.8; bigeminy: 1948.2 ± 298.7 mmHg.s^-1^), and a significant decrease in LV work (SR: 8834.4 ± 1214.0; bigeminy: 7291.0 ± 1068.) and dP/dt min (SR: -936.0 ± 177.8; bigeminy: -1462.3 ± 245 mmHG.s^-1^) (Figure 4E-I). Interestingly, premature extrasystole during bigeminy led to premature contractions that, on average, have a DP of only 50% of that of SR (Figure 4J).

**Figure 4.**
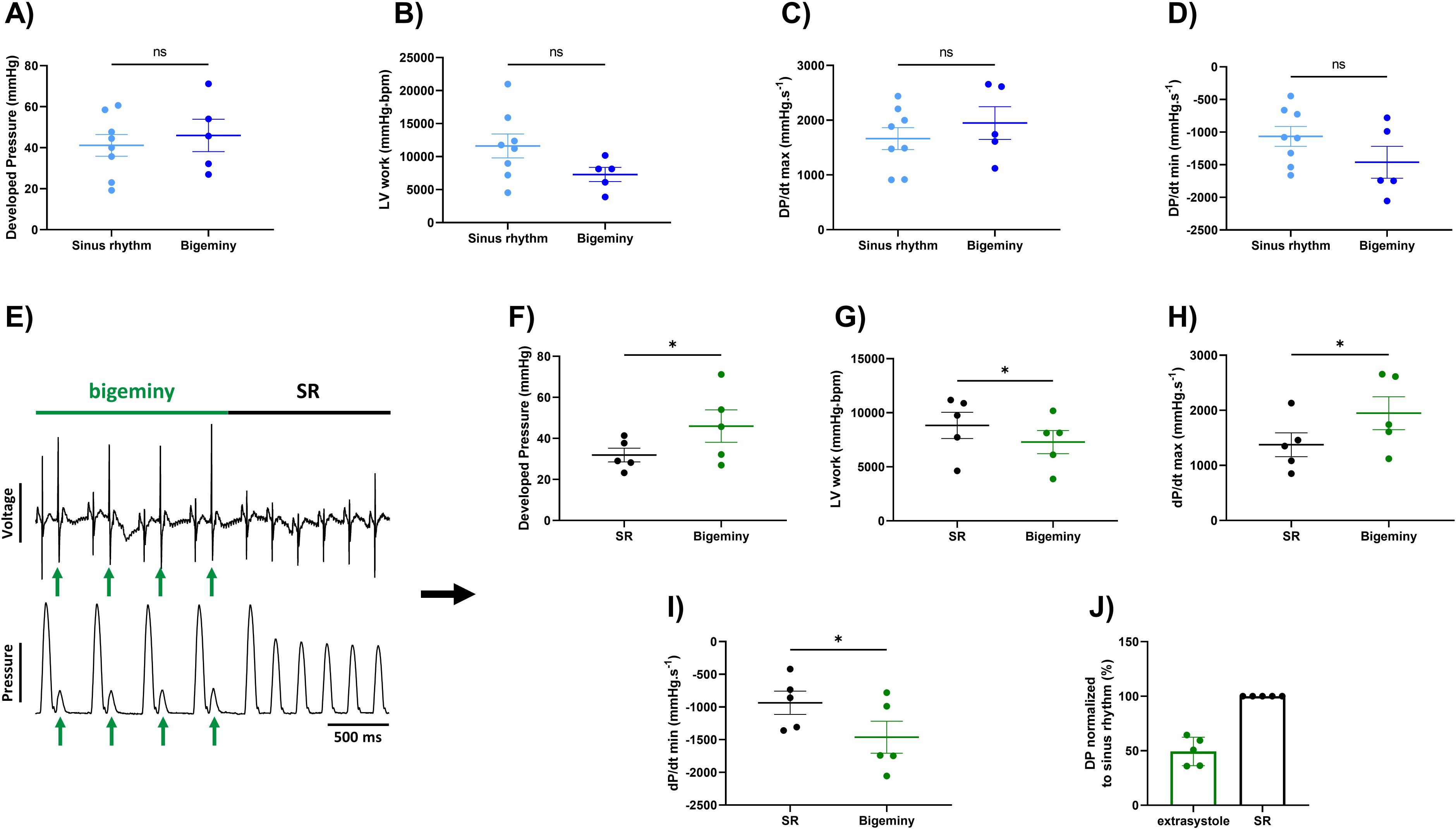
Altered mechanical function during episodes of bigeminy. Average developed pressure (A), left ventricular (LV) work (B), and maximal velocity of contraction (dP/dt max) (C) and relaxation (dP/dt min) (D) among hearts presenting with sinus rhythm or bigeminy at baseline (N=13). (E) Example trace recording of ECG (top) and left ventricular pressure (bottom) from one of the individual hearts showing a brief episode of bigeminy (green bar) followed by sinus rhythm (black bar). Green arrows indicate the extrasystoles. Average developed pressure (F), left ventricular (LV) work (G), and maximal velocity of contraction (dP/dt max) (H) and relaxation (dP/dt min) (I) between episodes of sinus rhythm (black) or bigeminy (green) in the same hearts (N=5). (J) Average developed pressure (DP) of the extrasystoles (green) normalized to average DP during episodes of sinus rhythm (SR, black) in the same hearts (N=5). *ns = non-significant, * p < 0.05, ** p < 0.01 with unpaired (A-D) and paired (F-I) t-test*.

### 4.3 Global ischemia-reperfusion decreased mechanical function, while electrical function remained mainly unchanged

This combined electromechanical assessment approach in *ex-vivo* Langendorff perfused hearts can be combined with any experimental conditions. Here we decided to perform global ischemia for 25 minutes followed by 1h of reperfusion. After ischemia, the mechanical function dropped drastically with a significant decrease in DP (baseline: 37.7 ± 4; reperfusion 10 min: 11.8 ± 2.4 mmHg), LV work (baseline: 9351 ± 1174; reperfusion 10 min: 2927 ± 545 mmHg*bpm), dP/dt max (baseline: 1509.8 ± 137.1; reperfusion 10 min: 551.5 ± 99.6 mmHg.s^-1^) and dP/dt min (baseline: -1005 ± 101.5; reperfusion 10 min: -344.3 ± 59.2 mmHg.s^-1^) at 10 minutes reperfusion (Figure 5A-E). Then, the mechanical function slowly recovered to baseline level with an average recovery of 79.3 ± 7.2 % for DP, 67.0 ± 9.8 % for LV work, 98.3 ± 11.0 % for dP/dt max, and 75.9 ± 8 % for dP/dt min at 60 minutes reperfusion (Figure 5A, Supplementary Figure 3A). Of note, the average coronary flow remained the same throughout the experiment, e.g., in the baseline perfusion and the reperfusion after global ischemia (Supplementary Figure 3B). In contrast to the ventricular function, electrical function stayed relatively stable with no significant change in heart rate, PR and QT interval, intra-atrial and intra-ventricular conduction times (Figure 5F-I, Supplementary Figure 3C, D). However, the ventricular effective refractory period was significantly increased after global ischemia (Figure 5J), indicating changes in cardiac repolarization. Besides, though not significant, heart rate tended to be more irregular after global ischemia-reperfusion, as shown by the increase of the average coefficient of variation (standard deviation/mean) of the inter-beat time (Supplementary Figure 3E).

**Figure 5.**
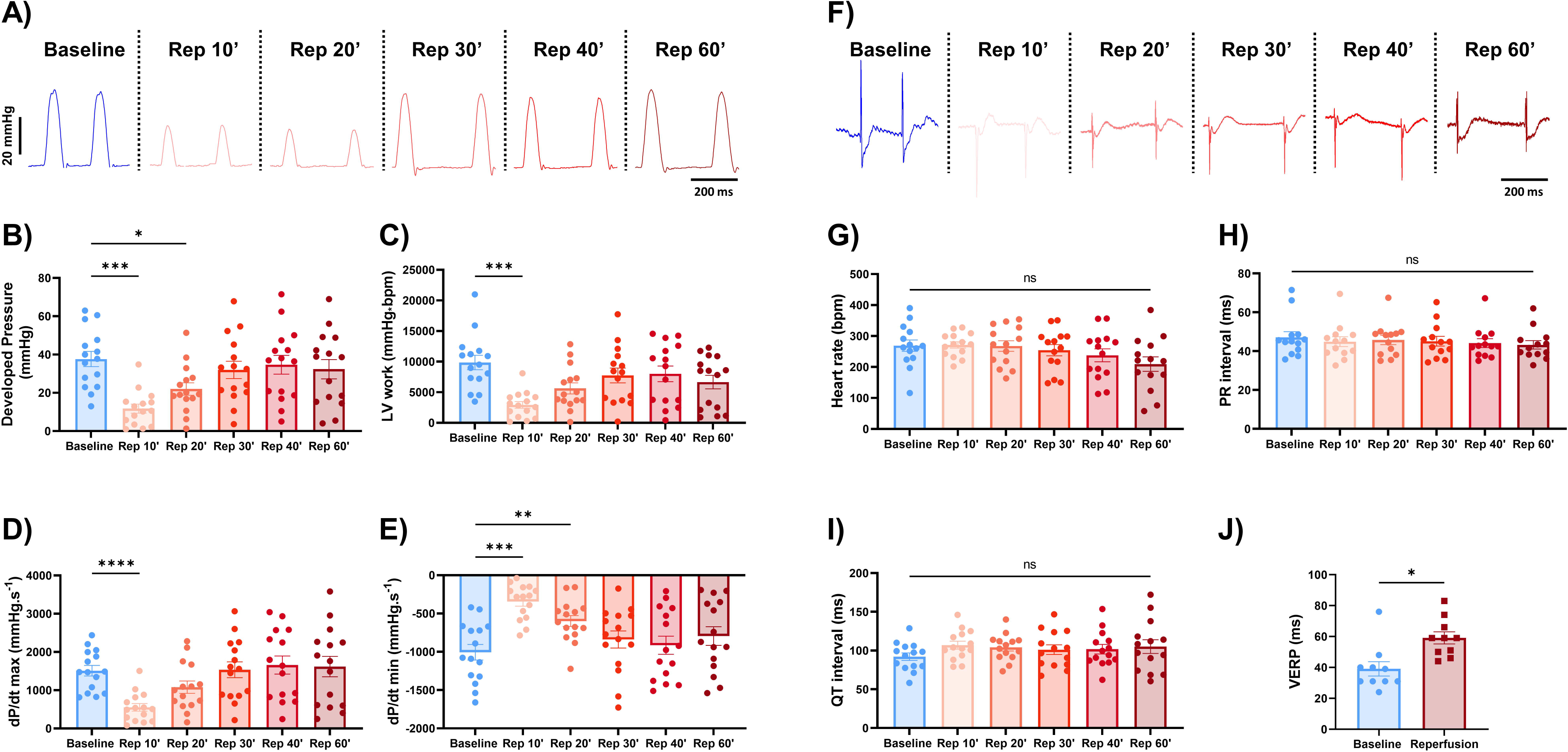
Global ischemia-reperfusion decreased mechanical function, while electrical function remained largely unaffected. (A) Example trace recordings of the left ventricular pressure and average developed pressure (B), left ventricular (LV) work (C) and maximal velocity of contraction (dP/dt max) (D) and relaxation (dP/dt min) (E) at baseline and 10, 20, 30, 40, and 60 minutes of reperfusion (Rep 10’, 20’, 30’, 40’, 60’) (N=15). (F) Example ECG recordings and average heart rate (G), PR interval (H) and QT interval (I) at baseline and 10, 20, 30, 40, and 60 minutes of reperfusion (Rep 10’, 20’, 30’, 40’, 60’) (N=15). (J) Average ventricular effective refractory period (VERP) between baseline (blue) and reperfusion (red) (N=10). *ns = non-significant, * p < 0.05, ** p < 0.01, *** p < 0.001, **** p < 0.0001 with one-way ANOVA followed by Dunnett’s multiple comparison test (compared to baseline) (B-E, G-I) or paired t-test (J)*.

Taken together, these results indicate that the ischemia-induced decrease in mechanical function is not due to or linked to changes in electrical function.

### 4.4 Global ischemia-reperfusion does not affect the electro-mechanical interaction

We then wondered whether global ischemia affects the electro-mechanical interaction observed at baseline. At the end of reperfusion, 12 hearts were in sinus rhythm, 1 showed sustained 1^st^ degree AV block, 3 had brief episodes of bigeminy accounting for less than 20% of the total time period (Figure 6A) and another one showed a brief episode of idioventricular rhythm faster than sinoatrial rhythm that was not observed at baseline (Figure 6A-C). Of note, apart from one heart in AV block, arrhythmic hearts at baseline and at the end of reperfusion were not the same, meaning that some hearts showed new arrhythmic behavior after global ischemia while others lost their arrhythmic pattern (Figures 3E and 6A). We again compared the mechanical function during episodes of bigeminy in the hearts presenting with this arrhythmia during reperfusion with that of sinus rhythm hearts (Supplementary Figure 4A-D). There was no significant difference in DP, LV work, dP/dt max and min. However, when we compared the mechanical function during episodes of bigeminy and sinus rhythm in the same hearts, there were increases in DP (SR: 40.5 ± 9.9; bigeminy: 50.8 ± 6.6 mmHg) and dP/dt max (SR: 1761 ± 337; bigeminy: 2222 ± 95.9 mmHg.s^-1^) and decreases in LV work (SR: 7968 ± 1158; bigeminy: 7098 ± 812 mmHg*bpm) and dP/dt min (SR: -1089 ± 197; bigeminy: -1451 ± 280 mmHg.s^-1^) (Figure 6E-F, Supplementary Figure 4E-F). In addition, the premature beats had on average a DP of 50% of that of the SR (Figure 6G). Therefore, the mechanical response to bigeminy during reperfusion is the same as what was observed at baseline (Figure 4F-J).

**Figure 6.**
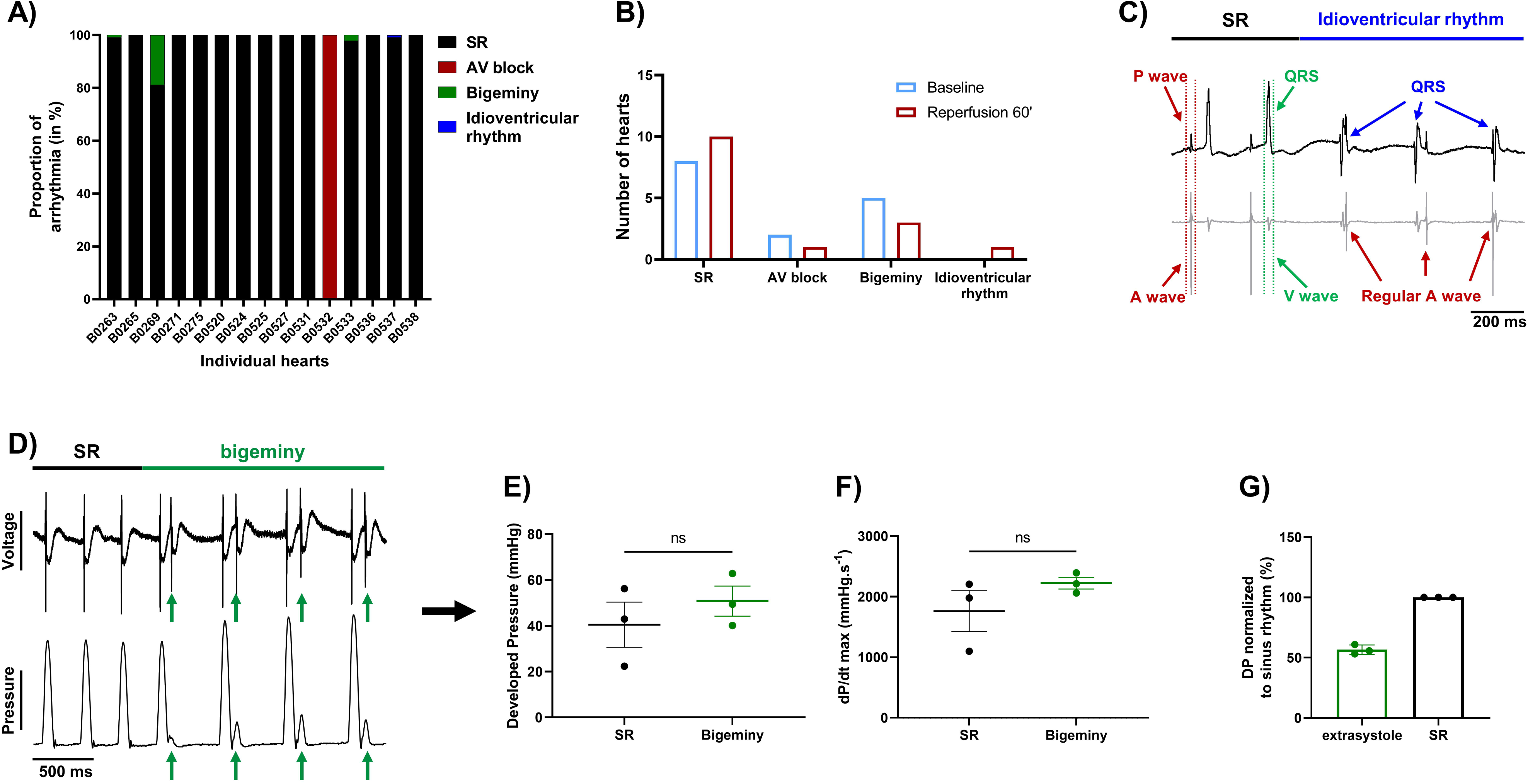
Effect of bigeminy episodes on mechanical function is preserved after global ischemia-reperfusion. (A) Proportion of arrhythmia occurrence for a 2-minute period in individual hearts at the end of reperfusion. Black = sinus rhythm (SR), red = AV block, green = bigeminy and blue = idioventricular rhythm (N=15). (B) Proportion of *ex-vivo* hearts showing sinus rhythm (SR), AV block, bigeminy or idioventricular rhythm at baseline (blue bars) and 60 minutes reperfusion (red bars) (N=15). (C) Example trace recording of ECG (top) and one signal trace (channel 6, see Figure 2) from the octapolar catheter (bottom, gray) from the individual heart showing sinus rhythm (SR, black bar) followed by an episode of idioventricular rhythm faster than the sinoatrial node rate (idioventricular rhythm, blue bar). Red and green arrows indicate atrial and ventricular signals, respectively, while blue arrows indicate QRS complexes during idioventricular rhythm. (D) Example trace recording of ECG (top) and left ventricular pressure (bottom) from one heart showing sinus rhythm (SR, black bar) followed by an episode of bigeminy (green bar) during reperfusion. Green arrows indicate the extrasystoles. Average developed pressure (E) and maximal velocity of contraction (dP/dt max) (F) between episodes of sinus rhythm (black) or bigeminy (green) in the same hearts (N=3). Average developed pressure (DP) of the extrasystoles (green) normalized to average DP during episodes of sinus rhythm (SR, black) in the same hearts (N=3). *ns = non-significant with paired t-test*.

These results suggest that the electro-mechanical interaction is preserved after global ischemia-reperfusion and that the decrease in mechanical function (Figure 5A) after ischemia is rather due to an ischemia-mediated decrease in intrinsic mechanical properties.

## 5 Discussion

This manuscript presents an approach to simultaneously assess electrical and mechanical function in *ex-vivo* perfused mouse hearts. We also demonstrate that this approach can be combined with varied experimental conditions such as global ischemia-reperfusion. In addition, although this Langendorff setup was designed for mouse hearts, this should be readily transposable to larger mammal hearts such as rabbits with some adjustments for size.

### 5.1 Simultaneous recording of cardiac mechanical and electrical function

Though there is a growing need for new approaches to study cardiac electro-mechanical interactions, most of the time, the contractile and electrical function of isolated *ex-vivo* perfused hearts are investigated separately (1). In this manuscript, we showed that these can be assessed simultaneously with a balloon inserted in the left ventricle combined with ECG electrodes and an intracardiac octapolar catheter inserted in the right atrium and ventricle (Figure 1,2). Classical ECG parameters (heart rate, PR, and QT interval) can be measured alongside left ventricular mechanical properties (DP, LV work, maximal velocity of contraction and relaxation) (Figure 2). With the addition of the octapolar catheter, electrical conduction properties with intra-atrial and intra-ventricular conduction times can also be measured (Figure 2). This can be further refined by stimulating the heart to obtain data on ventricular refractoriness. Overall, our results at baseline are similar to what was previously described in terms of heart rate, PR interval, QT interval, and VERP in the mouse heart (16), confirming that our modified Langendorff setup does not impact cardiac electrical function. Regarding mechanical function, the left ventricular developed pressure described in literature ranges from 40 to >120 mmHg (17–20) with the DP obtained in our isolated mouse hearts (37.7 ± 4 mmHg) ranking amongst the lower values. This could be explained by the fact that we used a perfusion pressure of 60 mmHg which is associated with lower DP values (18). In addition, in order to evaluate electrical and mechanical function relationships over a wide range, unlike most studies no exclusion criteria based on heart function were applied (21), which may account for both the relatively low DP values and high variability (Figure 2).

Interestingly, we found that the developed pressure was not dependent on the heart rate, nor was the relaxation rate dependent on the QT interval (Supplementary Figure 1). We also found that the QT interval was negatively correlated to the heart rate, which is in line with the rate-dependence of the QT interval known in humans and larger mammals (22,23). However, the QT dependence of heart rate in mice is matter of debate, with studies showing either a dependence or no dependence in unanesthetized mouse hearts (24–26), which resulted in the lack of an accurate QT correction formula for mice.

### 5.2 Arrhythmia and subsequent mechanical reaction

The advantage of combining ECG recordings and the intracardiac octapolar catheter is that we can better define episodes of arrhythmia (27,28). For example, we found that almost half of the hearts, at baseline, had isolated extrasystoles or brief episodes of bigeminy (Figure 3, Supplementary Figure 2). These extrasystoles can originate from the atria (PAC) or the ventricles (PVC), which can easily be seen on the octapolar catheter signals with an extra A or V wave. Although this was not investigated, we could also obtain a rough idea of the origin of the ectopic signal based on which octapolar electrode is reached first. Therefore, the octapolar catheter is a real asset to study regional differences in the right atrium and ventricle, especially regarding blocks of conduction that might not be spotted on the one-lead pseudo-ECG.

Here we investigated the effect of brief episodes of bigeminy on the cardiac mechanical function. Comparison between episodes of bigeminy and episodes of sinus rhythm within the same hearts showed a significant improvement in DP and maximal velocity of contraction and relaxation in the subsequent beats following the extrasystole (Figure 4I-J). In contrast, the contraction triggered by the extrasystole was only 50% of those during sinus rhythm (Figure 4J), which is consistent with the phenomenon of “pulse deficit” observed in bigeminy patients: due to reduced diastolic filling and a short coupling interval of the extrasystole, there is a 2:1 pulse deficit in these patients that leads to a regular but slower pulse rate (29). Given the reduced contraction observed during the extrasystole in our heart models, we can assume that these would lead to insufficient ejection fraction and to the same 2:1 pulse deficit *in-vivo*. Hence, we showed here that our approach to simultaneously assessing electro-mechanical function can visualize recognized pathophysiological mechanisms and could be used to study the electro-mechanical interaction on a beat-to-beat scale. Indeed, with our modified setup, it is possible to see how each beat behaves electrically and mechanically. One can then study how a change in the electrical signal (arrhythmia, use of pharmacological agents targeting ion channels, pacing) can affect the mechanical function. Inversely, one can see how changes in the mechanical function (change in pressure/volume, use of pharmacological agents targeting the contractile proteins) can influence electrical properties.

### 5.3 Global ischemia-reperfusion effect on electro-mechanical interaction

As mentioned, this combined electro-mechanical assessment approach can be used for any experimental conditions on *ex-vivo* Langendorff-perfused hearts. Here we showed an example of global ischemia-reperfusion with continuous monitoring of both electrical and mechanical function. Interestingly, except for the increased VERP, global ischemia-reperfusion did not affect electrical parameters, even at an early stage of reperfusion (10 minutes) (Figure 5F-J, Supplementary Figure 3C-E). On the other hand, the mechanical function was decreased after ischemia and slowly recovered at the end of reperfusion (Figure 5A-E). This decrease in mechanical function after ischemia-reperfusion is referred to as “myocardial stunning” (30). It is most likely due to an increase in reactive oxygen species (ROS) and alterations in excitation-contraction coupling. Indeed, ROS generation after ischemia impairs sarcoplasmic reticulum function and provokes oxidative modifications of myofibrillar proteins (31–33). Upon post-ischemic reperfusion, cardiomyocytes also suffered from cytosolic Ca^2+^ overload due to increased Na^+^/H^+^ exchange and subsequent reverse mode of the Na^+^/Ca^2+^ exchanger (32,34,35), which contributes to ROS generation and, conversely, is aggravated by the ROS-induced impairment of sarcoplasmic reticulum function. In the end, oxidative modification of myofibrillar proteins leads to a decrease in Ca^2+^ responsiveness where Ca^2+^ dynamics (Ca^2+^ transient generation) is normal, but the contractile response is reduced (32,36,37). This is in line with our results, which show an unaffected electrical activity during reperfusion while the contractile response is decreased. However, we cannot rule out the possibility that part of this decrease in mechanical heart function may also be due to lethal reperfusion injury (8).

Interestingly, during reperfusion, we still observed the same contractile response during bigeminy as what was previously seen at baseline. Indeed, the extrasystole-triggered contractions were only 50% as strong as in sinus rhythm, while the subsequent contraction had an increased DP and maximal velocity of contraction and relaxation (Figure 6D-G, Supplementary Figure 4E-F). Therefore, the electro-mechanical interaction was preserved after ischemia-reperfusion, despite decreased mechanical function. This means that our new approach for assessing the electro-mechanical interaction could also be adapted to investigate the cardioprotective effect of a drug on ischemia-reperfusion injury as well as their effect on excitation-contraction and mechanical-electrical feedback.

## 6 Limitations

Our approach was designed for heart perfusion with the Langendorff method, a technique for which the perfusion is forced into the coronaries with little to no filling of the cardiac chambers. Hence, the ventricles are not exposed to blood/perfusate-induced mechanical strains. Therefore, it will be interesting to try adapting our approach to a perfusion system of working hearts to recreate a more physiological electro-mechanical interaction.

## 7 Conclusion

Our study presents an approach for simultaneously assessing electrical and mechanical function in Langendorff-perfused mouse hearts on a beat-to-beat scale. We show that this approach can be combined with any experimental conditions in Langendorff-perfused heart with an example of global ischemia-reperfusion. With this approach, we hope to better understand the complex mutual influence of cardiac electrical and mechanical function under physiological and pathophysiological conditions.

## Author contributions

SL, KEO and YD conceived the project. JL, RO, AS, AO, ME, MB, SN and NA did the experiments and acquired the data. JL and RO analyzed the data and drafted the manuscript. BMS and MS created and took care of the mouse strain line. All authors revised the manuscript, approved the final version and agreed to be accountable for all aspects of the work.

## Funding

This study was supported by the SNSF through a NRP 78 “Covid-19” Grant (CoVasc, No. 4078P0_198297) to YD, KEO and SL.

## Acknowledgements

The authors thank Maria Essers for her technical support regarding the isolated mouse heart system.

## Conflict of interest

The authors declare that the research was conducted in the absence of any commercial or financial relationships that could be construed as a potential conflict of interest.

## Notes

### Competing Interest Statement

The authors have declared no competing interest.

