## Supplementary figures for "Simultaneous assessment of mechanical and electrical function in Langendorff-perfused ex-vivo mouse heart"

### *Supplementary Material*

**Supplementary Figure 1**

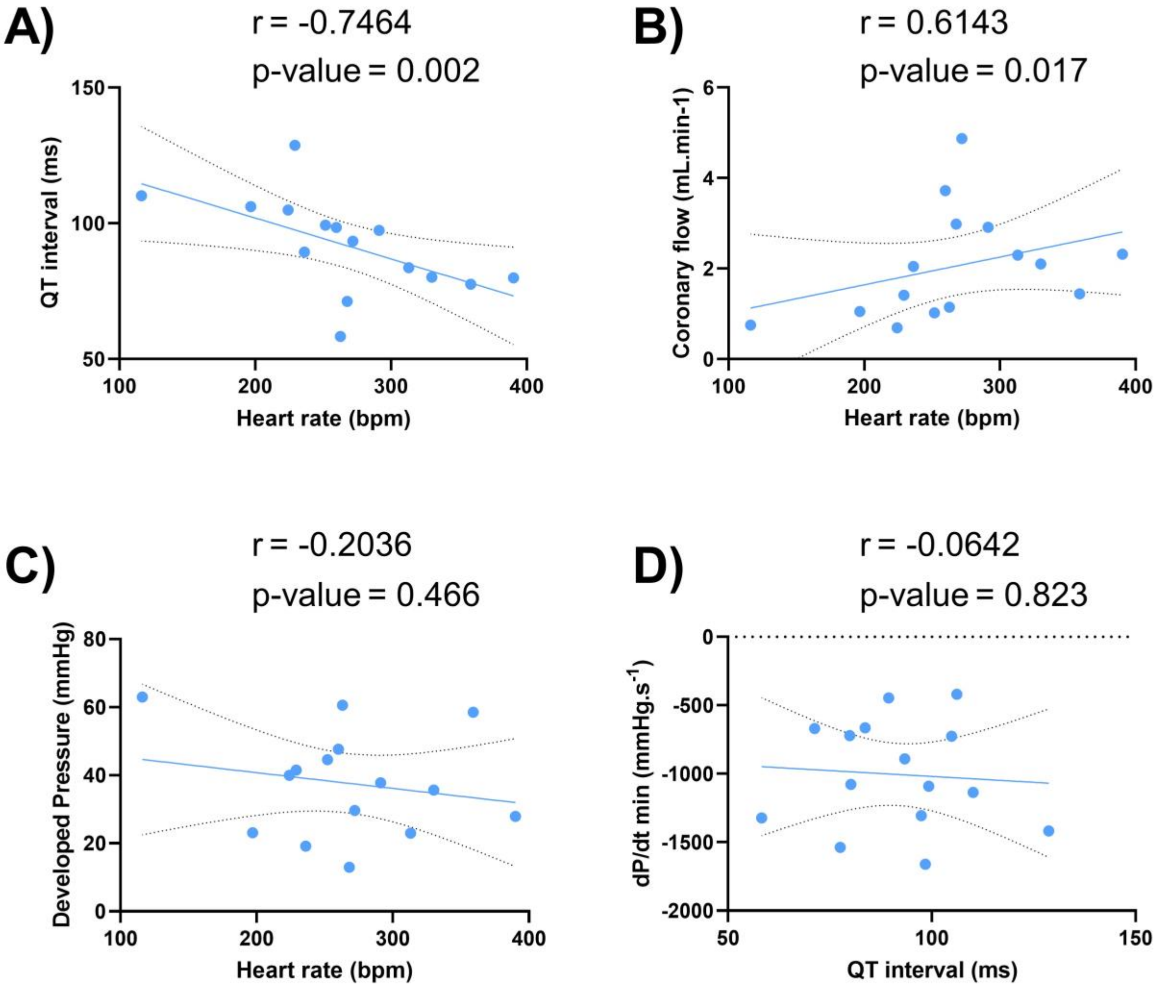


**Supplementary Figure 1. Correlation between heart rate and electrical / mechanical function.** Linear regression of the correlation between QT interval and heart rate (A), coronary flow and heart rate (B), developed pressure and heart rate (C), and maximal velocity of relaxation (dP/dt min) and QT interval (D) at baseline (N=15). *Statistical significance is reached when p-value < 0.05 with a Spearman correlation test.*

**Supplementary Figure 2**

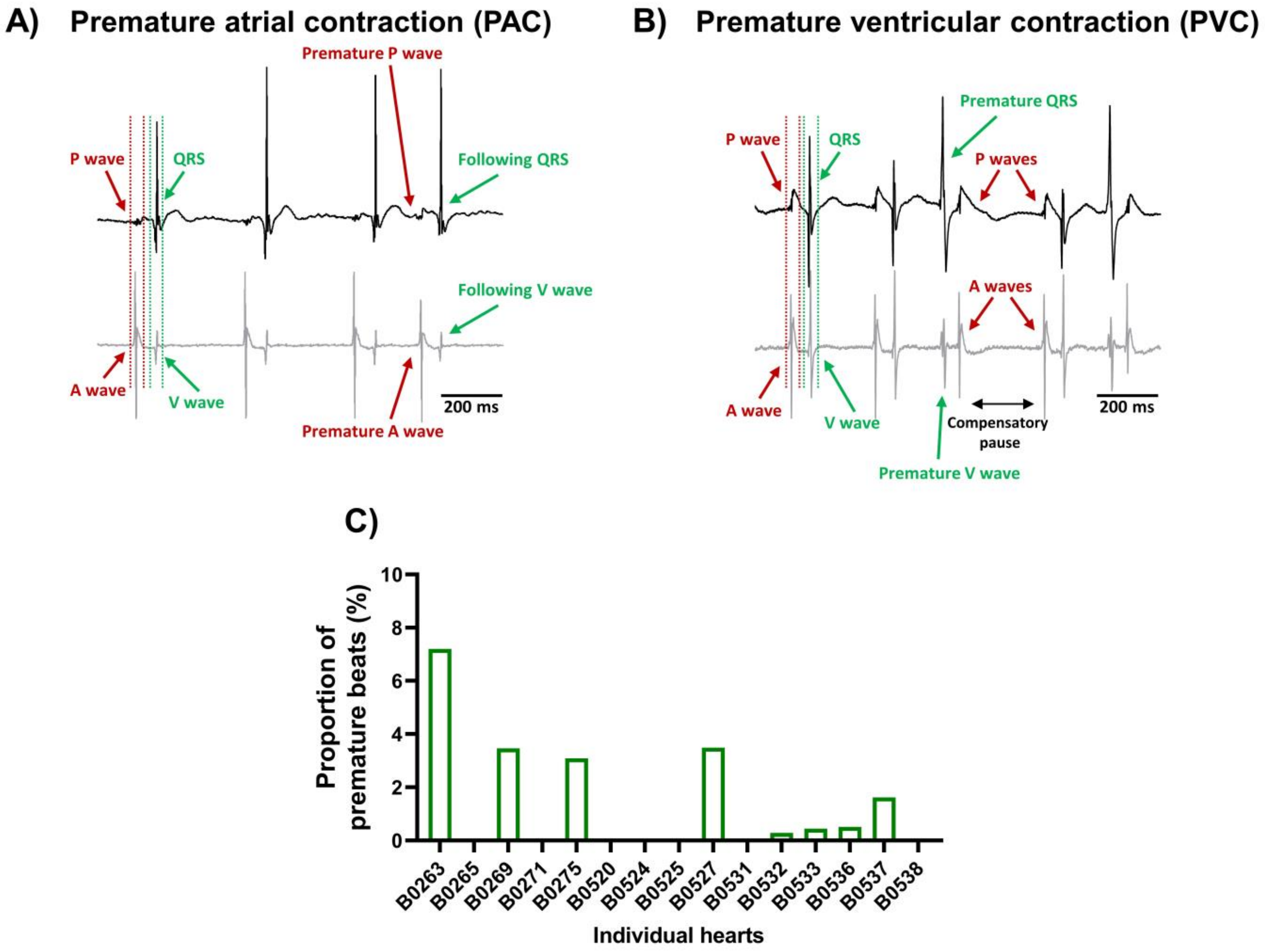


**Supplementary Figure 2. Spontaneous isolated premature beats at baseline.** Example recordings of the electrocardiogram trace (top, black) and one signal trace (channel 6, see Figure 2) from the octapolar catheter (bottom, gray) of a premature atrial contraction (PAC) (A) and a premature ventricular contraction (PVC) (B) at baseline. Red and green arrows indicate atrial and ventricular signals, respectively. (C) Proportion of premature beats normalized to the total number of beats during the considered time-window at baseline.

**Supplementary Fig. 3**

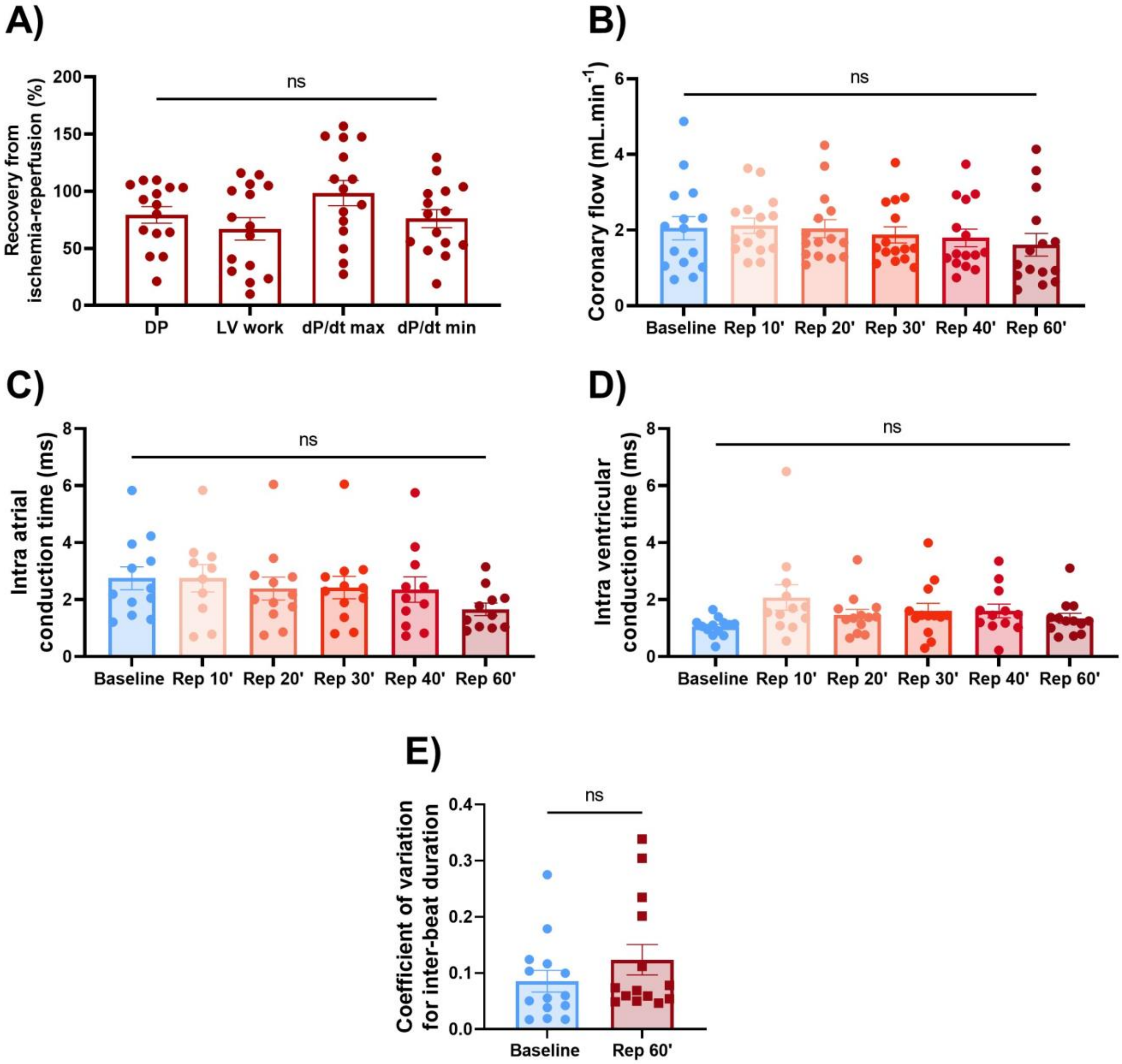


**Supplementary Figure 3. Effects of ischemia-reperfusion on mechanical and electrical function.** (A) Average percentage recovery from ischemia at the end of reperfusion normalized to baseline values for developed pressure, LV work and maximal velocity of contraction (dp/dt max) and relaxation (dP/dt min) (N=15). Average coronary flow (B) and atrial (C) and ventricular (D) conduction times at baseline and 10, 20, 30, 40, and 60 minutes of reperfusion (Rep 10', 20', 30', 40', 60') (N=15) (E) Average coefficient of variation for inter-beat duration between baseline and 60 minutes of reperfusion (Rep 60'). *ns = non-significant with one-way ANOVA followed by Dunnett's multiple comparison test (A-D) or non-parametric paired t-test (Wilcoxon test) (E).*

**Supplementary Fig. 4**

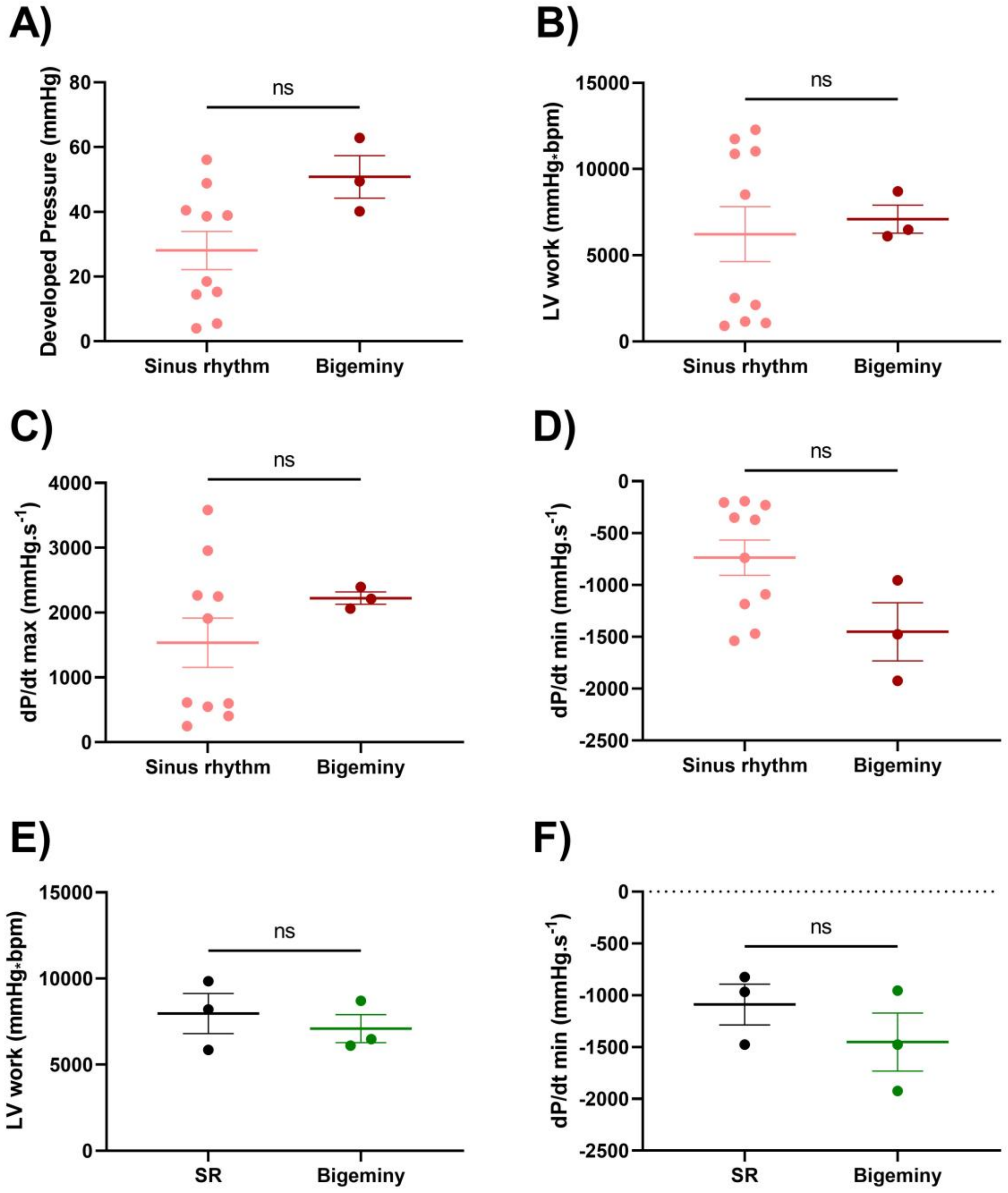


**Supplementary Figure 4. Effects of arrhythmia on mechanical function after global ischemia-reperfusion.** Average developed pressure (A), left ventricular (LV) work (B), and maximal velocity of contraction (dP/dt max) (C) and relaxation (dP/dt min) (D) between individual hearts presenting with sinus rhythm or bigeminy at 60 minutes reperfusion (N=13). Average LV work (E) and maximal velocity of relaxation (F) between episodes of sinus rhythm (black) or bigeminy (green) in the same hearts (N=3). *ns = non-significant with unpaired (A-D) or paired (E-F) t-test.*
